# No effect of triple-pulse TMS medial to intraparietal sulcus on online correction for target perturbations during goal-directed hand and foot reaches

**DOI:** 10.1101/645853

**Authors:** Daniel S. Marigold, Kim Lajoie, Tobias Heed

## Abstract

Posterior parietal cortex (PPC) is central to sensorimotor processing for goal-directed hand and foot movements. Yet, the specific role of PPC subregions in these functions is not clear. Previous human neuroimaging and transcranial magnetic stimulation (TMS) work has suggested that PPC lateral to the intraparietal sulcus (IPS) is involved in directing the arm, shaping the hand, and correcting both finger-shaping and hand trajectory during movement. The lateral localization of these functions agrees with the comparably lateral position of the hand and fingers within the motor and somatosensory homunculi along the central sulcus; this might suggest that, in analogy, (goal-directed) foot movements would be mediated by medial portions of PPC. However, foot movement planning activates similar regions for both hand and foot movement along the caudal-to-rostral axis of PPC, with some effector-specificity evident only rostrally, near the central regions of sensorimotor cortex. Here, we attempted to test the causal involvement of PPC regions medial to IPS in hand and foot reaching as well as online correction evoked by target displacement. Participants made hand and foot reaches towards identical visual targets. Sometimes, the target changed position 100-117 ms into the movement. We disturbed cortical processing over four positions medial to IPS with three pulses of TMS separated by 40 ms, both during trials with and without target displacement. We timed TMS to disrupt reach execution and online correction. TMS did not affect endpoint error, endpoint variability, or reach trajectories for hand or foot. While these negative results await replication with different TMS timing and parameters, we conclude that regions medial to IPS are involved in planning, rather than execution and online control, of goal-directed limb movements.

## Introduction

The dominant view of parietal organization is that specialized posterior parietal cortex (PPC) sub-regions are responsible for the planning of movements involving specific effectors. This view is based primarily on research in non-human primates. For example, monkey lateral intraparietal region is mainly involved in saccade planning, the neighboring parietal reach and medial intraparietal regions in hand reach planning, and the anterior intraparietal region in grasp planning [1,2]. In humans, this effector specificity is less clear: functional magnetic resonance imaging (fMRI) activity in PPC for eye and hand planning largely overlaps, though with some biases for one or the other effector in certain regions [2,3–5]. Only a few studies have compared hand and foot processing in PPC and found surprisingly large overlap for the two effectors [4–8]. Specifically, posterior regions involved in hand reaching (e.g., superior parieto-occipital cortex, SPOC) and a region in the superior parietal lobule (SPL) appear just as actively involved in foot as in hand movements, and only regions directly neighboring effector-specific sensorimotor cortex show clear effector-specificity during motor planning. These findings challenge the idea that effector-specificity is a guiding principle of PPC organization and suggest that PPC may, instead, be organized according to functional aspects (such as the online monitoring and correcting of movement trajectories towards target objects and locations) rather than body parts.

Rapid changes in limb trajectory due to changes in a target’s location are a necessity given the dynamic nature of the world we live in. To accomplish such online corrections of goal-directed motor acts the brain must continually monitor limb state. This state estimate is derived from a combination of incoming sensory input and internally generated predictions about upcoming sensory feedback, the latter of which is presumably based on a forward internal model that uses a copy of the motor command and knowledge of limb dynamics [9]. When target location changes, the estimate of limb state can be compared to the estimate of the new location, and the difference can be used to create a new motor command to produce a change in limb trajectory.

PPC may encode the current state estimate [10,11] and play a key role in computing the distance (or motor error) between effector and target locations [2,12–16]. In support, neurons in the parietal reach region were found to encode the current movement angle of a cursor that monkeys controlled via a joystick to make reaches to targets [10]. In addition, neuroimaging studies suggest that the anterior intraparietal sulcus is active in the early phase of adapting to altered visual input when reaching error is high [17,18]. Furthermore, trial-by-trial correlation of fMRI activation in this region and hand reaching error suggests that this region is involved in error processing [18]. Thus, disruption of the PPC after movement onset should impair or prevent online corrections to limb trajectory.

Single-cell recordings demonstrate that discharge activity of area 5 PPC neurons is modified when online corrections of reaches (in monkeys) or gait (in cats) are made [19–21]. Moreover, lesions to the human PPC can impair the ability to correct the trajectory of a reach when the target is unexpectedly moved to a new location during the movement [22]. Using transcranial magnetic stimulation (TMS), Desmurget et al. [13] found that perturbation applied to medial intraparietal sulcus at the onset of a goal-directed reach disrupted trajectory corrections after unexpected target shifts. Similarly, TMS to anterior intraparietal sulcus impairs the ability to produce the appropriate forearm orientation when the grasp object is suddenly rotated [23] and reduces the ability to online correct to changes in visual feedback of a target or the hand during reaches [24]. The fact that disruption to these regions affects grasping, forearm positioning, and hand localization has led these researchers to suggest that the involvement of human anterior intraparietal sulcus in online corrections is effector-independent.

Previous work that has addressed the question of effector-specificity (beyond a comparison of hand and eye movement) has investigated simple motor tasks and used fMRI. However, fMRI is inherently correlative in nature. In contrast, TMS can temporarily disrupt neural activity in a focal brain area and thereby test for the causal role of the specific candidate region in the neural process under scrutiny [25]. Here, we sought to gain causal evidence for effector-specific versus functional (i.e., effector-independent) involvement of PPC regions in the planning and control of online corrective hand and foot movements.

## Materials and methods

To test if PPC is organized in an effector-specific manner, participants performed goal-directed hand and foot movements. During movement execution the target occasionally changed (or ‘jumped’) to a new location, forcing participants to adjust their movement to maintain accuracy. We applied TMS to regions identified with fMRI as active specifically for only one effector, or for both hand and foot movement planning. This tests if the ability to make online corrective movements during goal-directed hand and foot reaches is reduced when the function of specific regions of the PPC are disrupted. TMS to regions involved in reach correction of a specific effector should impair reach adjustment to target jumps for the respective effector, but not for others. TMS to regions involved in reach correction of all effectors should, in contrast, impair reach adjustment to target jumps independent of the presently used effector. Finally, TMS to regions that are not involved in mediating online correction will not result in modulation of reach trajectory for any effector. This approach allows us to dissociate whether the targeted regions are effector-specific, or whether they support specific, sensorimotor-related functionality for all (tested) effectors.

All data used to produce statistical results and figures are available at https://osf.io/h3jym/.

### Participants

Eighteen individuals (aged 26.3 ± 3.7 years; 5 males, 13 females) with no known musculoskeletal, neurological, or visual disease participated in this study. Seventeen participants were right hand dominant and seventeen were right leg dominant. Hand dominance was reported by participants. Foot dominance was assessed by asking which leg a participant uses to kick a soccer ball. One participant did not have a dominant hand and one participant did not have a dominant leg. The ethics committee of the German Psychological Society (DGPs) approved the study, and all participants gave informed written consent before performing the experiments.

### Experimental setup

Participants performed hand and foot reaches to a target with their right limbs while sitting on a chair with a footrest. Participants placed their chin on a chin rest secured to the side of the chair to reduce movement of the head and facilitate TMS. A computer screen (48.0 × 29.6 cm; frame rate of 60 Hz) positioned in front displayed a circular target (diameter = 1.0 cm; light magenta) against a black background. Participants started each trial with either their right index finger or great toe on a trigger button on the chair. The foot and hand reach distance to the target screen varied between participants to ensure comfort (range = 29 to 31 cm) but remained equal for both limbs within a participant.

A touch screen mounted in front of the computer screen recorded the endpoint position of the hand and foot. We fixed the touch screen and chair in a rigid metal construction so that all distances remained constant, and the touchscreen was a non-moveable, wall-like target area. To determine hand and foot trajectories, a motion-capture system (Visualeyez VZ4000v, Phoenix Technologies Inc., Vancouver, Canada; sampling rate 100 Hz) positioned overhead recorded position markers placed on the right index finger, dorsal surface of the right hand midway along the second metacarpal bone, right great toe, and dorsal surface of the right foot midway along the first metatarsal bone.

### Location of brain sites and TMS protocol

We used frameless stereotaxic neuronavigation (BrainVoyager TMS Neuronavigator, Brain Innovation B.V., Maastricht, The Netherlands) to localize parietal brain sites for TMS and to monitor TMS coil position during the hand and foot reaches. Prior to testing, we acquired T1-weighted high-resolution structural MR images at 3T in a Siemens Trio 3T MR scanner (Siemens, Erlangen, Germany) to reconstruct the three-dimensional anatomy of each participant’s brain.

We selected four regions of the left PPC for TMS stimulation based on our previous work [4,5] and that of others [26–29]. We applied TMS to the target Talairach coordinate at the center of the selected regions. For each participant, we adjusted target coordinates to individual anatomy by reversely applying the transformation which warped the individual brain anatomy into Talairach space. If a given Talairach coordinate was outside the brain of an individual participant, we used the nearest alternative Talairach location instead.

The mean stimulated Talairach coordinates across our sample, illustrated in Fig. 1B, and were (from most posterior to most anterior):

**Fig 1.**
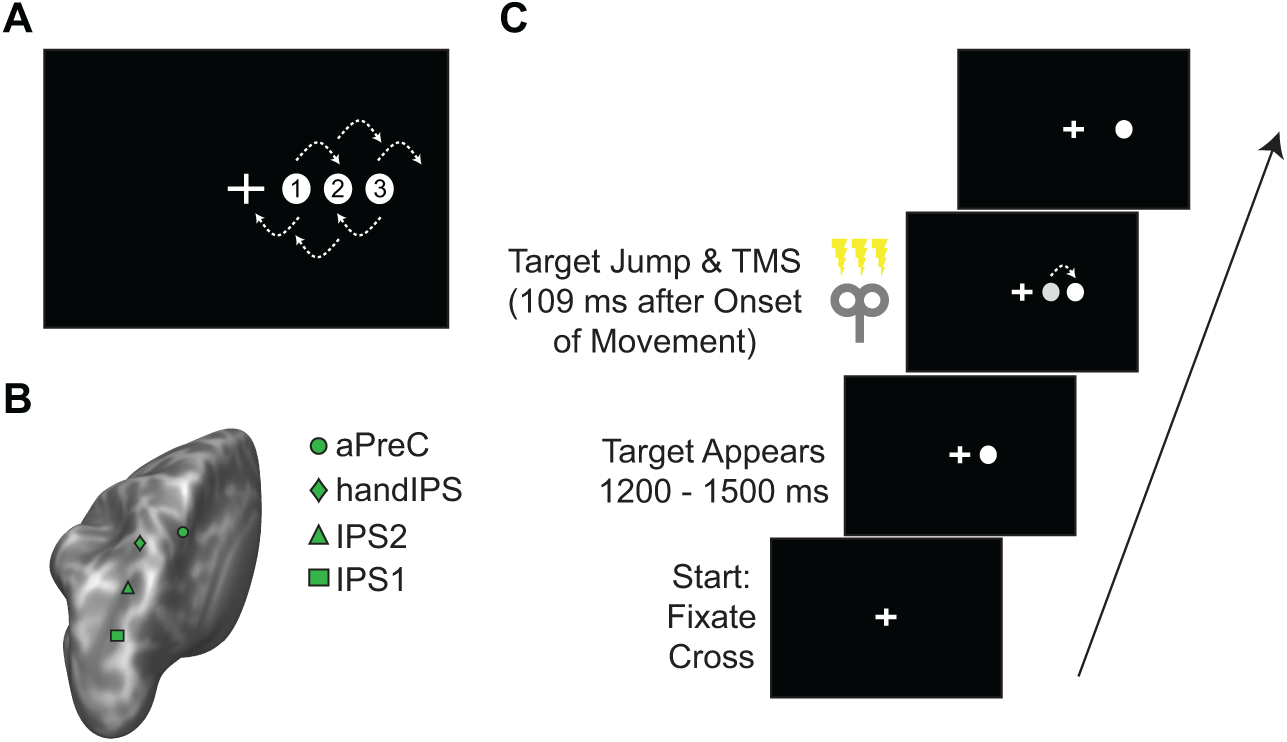
Experimental set-up and design. (A) A single target appeared in one of thre remained in the same position or either ‘jumped’ to the next position left or ri magnetic stimulation (TMS) to one of four posterior parietal cortex locations sulcus region (IPS1), a medial intraparietal sulcus region (IPS2), handIPS, and trial began with participants fixating a central cross. The target appeared at ran ‘jumped’ to a new location. In 50% of jump and non-jump trials, we applied tr

1. *x* = −22.0, *y* = −75.4, *z* = 38.3, nominally referred to hereafter as IPS1 (e.g., [4,28]), and also very close to a region referred to as superior parieto-occipital sulcus or cortex (sPOS, SPOC) in other work (e.g., [30–32]; see Heed et al. [4]). This region was previously found active for both hand and foot motor planning [4,5,7].
2. *x* = −19.0, *y* = −71.6, *z* = 46.9, nominally referred to hereafter as IPS2 (e.g., [28]). This region has been referred to as medial IPS in some studies [4,5] and was previously found active for both hand and foot motor planning [4,5,7].
3. *x* = −26.2, *y* = −56.4, *z* = 55.5, nominally referred to hereafter as handIPS. This region was previously found active for hand planning [5,33], involved in online corrections of reach-to-grasp actions [34,35], and lies between two hand-specific regions in our previous study that compared hand and foot reaching [7]. The region has previously been referred to as medial IPS [33], anterior IPS [5], and Brodmann area 7 [35].
4. *x* = −12.6, *y* = −48.8, *z* = 52.6, nominally referred to hereafter as anterior precuneus, or aPreC. This region was previously found active for foot, but not hand, motor planning [4,5,7].

We also conducted ‘sham’ conditions for movements of the foot and hand that we randomly interspersed between PPC conditions. In sham blocks, we held the edge of the TMS coil over the PPC to direct the induced magnetic field away from the brain [32,36]. We also applied test pulses to determine whether stimulation of each PPC site evoked a motor response in the hand or arm. This involved a single pulse to the specific site when the participants held their arms fully extended in front of them [36]. We found no evidence of a motor response during these test pulses in any participant.

During the experiment, we administered three pulses at 25 Hz 40 ms apart using a PowerMAG 100 TMS system (MAG & More GmbH, Munich, Germany) and an 8-cm figure-of-eight coil with the handle pointed backwards at an angle of 45° with respect to the anterior-posterior axis for all PPC stimulation sites. We delivered stimulation at 100% of individual resting motor threshold (similar to Striemer et al. [36] and between the 90% and 110% used by Reichenbach et al. [24] and Tunik et al. [23], respectively) of the right small hand muscles. We defined resting motor threshold of these muscles as the minimum stimulation intensity needed to evoke a visible twitch on 5 out of 10 trials when we placed the TMS coil over the hand area of the primary motor cortex (mean = 47.8 ± 6.1 % of stimulator output, range = 37 to 60.5 %). These stimulation parameters are in accordance with established safety standards [37].

### Experimental protocol

Initial target position varied pseudo-randomly from trial to trial between one of three different locations, 4.2, 8.4, or 12.6 cm to the right of midline (see Fig. 1A); all three locations were chosen with equal probability. Trial timing is illustrated in Fig. 1C. To begin a trial, participants placed either their index finger or great toe on a start button and fixated a central cross. A single reach target appeared randomly between 1,200 and 1,500 ms. Contact of the touch screen with the finger or toe extinguished the target. To initiate a subsequent trial the participant returned to the start button. If the participant started moving before presentation of the target, within 100 ms of target appearance, or if the movement started later than 1,500 ms after target presentation, we aborted the trial. We repeated these aborted trials at the end of the block. We also repeated a trial if the movement duration fell outside of 350 to 500 ms to ensure a relatively constant movement velocity. A message indicating that the movement was too slow or too fast appeared on the screen in these instances.

Each practice and experimental (i.e., PPC site) condition consisted of a block of 72 trials. In 36 of these trials (50%), the target ‘jumped’ shortly after foot or hand lift-off from the start button to a new location 4.2 cm left or right of its original position (i.e., jump trials). We specified for the jump to occur 100 ms after the lift-off; however, due to the refresh rate of the monitor, it could only be drawn every 16.6 ms, so that the jump occurred 100-117 ms (mean: 109 ms) after lift-off. Targets jumped left or right with equal probability. In the remaining 36 trials (50%), the target remained in the original position throughout the movement (i.e., non-jump trials).

We applied triple-pulse TMS in 18 of the non-jump and jump trials (50% of trials, respectively). In both types of trials, the first TMS pulse occurred together with the drawing of the first screen refresh following 100 ms after lift-off. Thus, in target jump trials, the first TMS pulse coincided with the target jump. In trials without target jump, the TMS pulses were timed identically, including temporal jitter related to screen-refresh.

Participants practiced the task for the first four blocks (two blocks with the foot and two blocks with the hand in alternating orders). The remaining ten blocks consisted of TMS applied to one of the four PPC sites (IPS1, IPS2, handIPS, aPreC) or sham TMS. We randomized the order of PPC/sham blocks. Furthermore, we rotated between hand and foot blocks, and we counterbalanced the order of the starting limb across participants. In total, each participant performed 144 practice and 360 valid experimental hand reaching trials, and equivalent numbers of foot reaching trials during the experiment.

### Data and statistical analyses

We analyzed data in MATLAB (The MathWorks, Natick, MA), and assessed statistical significance with JMP 13 software (SAS Institute Inc., Cary, NC). We performed separate analyses for each effector and brain site (or sham) block. To assess the accuracy of foot and hand reaches to the targets, we calculated endpoint error on each trial based on the Euclidean distance between the target center and position of the finger or toe on the touch screen. We calculated the area of 95% prediction ellipses fit to the endpoint errors [38]. In addition, we quantified mean movement times. We defined movement time as the interval between button release and contact with the target on the touch screen. To determine differences in error and movement time, we used two-way (Jump x TMS) ANOVAs with participant as a random effect for each limb (hand, foot) and brain site (IPS1, IPS2, handIPS, aPreC, Sham) separately. Because ellipse area did not follow a normal distribution, we log transformed the data before subjecting them to the ANOVAs. Our data revealed virtually no statistically significant effects. We therefore did not apply corrections for multiple comparisons.

To determine whether TMS affected the trajectory of online movement corrections to target jumps, we first filtered kinematic data using a 4^th^-order, zero-lag, Butterworth filter with a low-pass cutoff of 10 Hz. We used the vector position of the finger or toe marker for hand and foot reaches, respectively, and subsequently analyzed this trajectory data.

## Results

### General behavior

Participants performed goal-directed hand and foot movements, during which a target occasionally ‘jumped’ to a new location, forcing them to adjust their movement to maintain accuracy. We applied TMS to four different regions of the PPC that are known to be active during movement planning for either hand or foot, or for both. Participants initiated hand movements 313 ± 48 ms and foot movements 333 ± 52 ms after the appearance of a target in one of three positions to the right of midline.

Regardless of condition, participants corrected their hand and foot movements to the sudden change in target position (see Fig 2, for example). These online corrections occurred approximately 240 ms following the target jump. Movement times differed significantly between target-jump trials and non-jump trials (p < 0.0001, all brain site conditions). However, we found no effect of TMS on the movement times (non-significant TMS main effect and TMS x Jump interaction, p > 0.05 for all brain site conditions). Average movement time (pooled across all trials and brain sites) was 439 ± 27 ms for hand reaches and 443 ± 25 ms for foot reaches.

**Fig 2.**
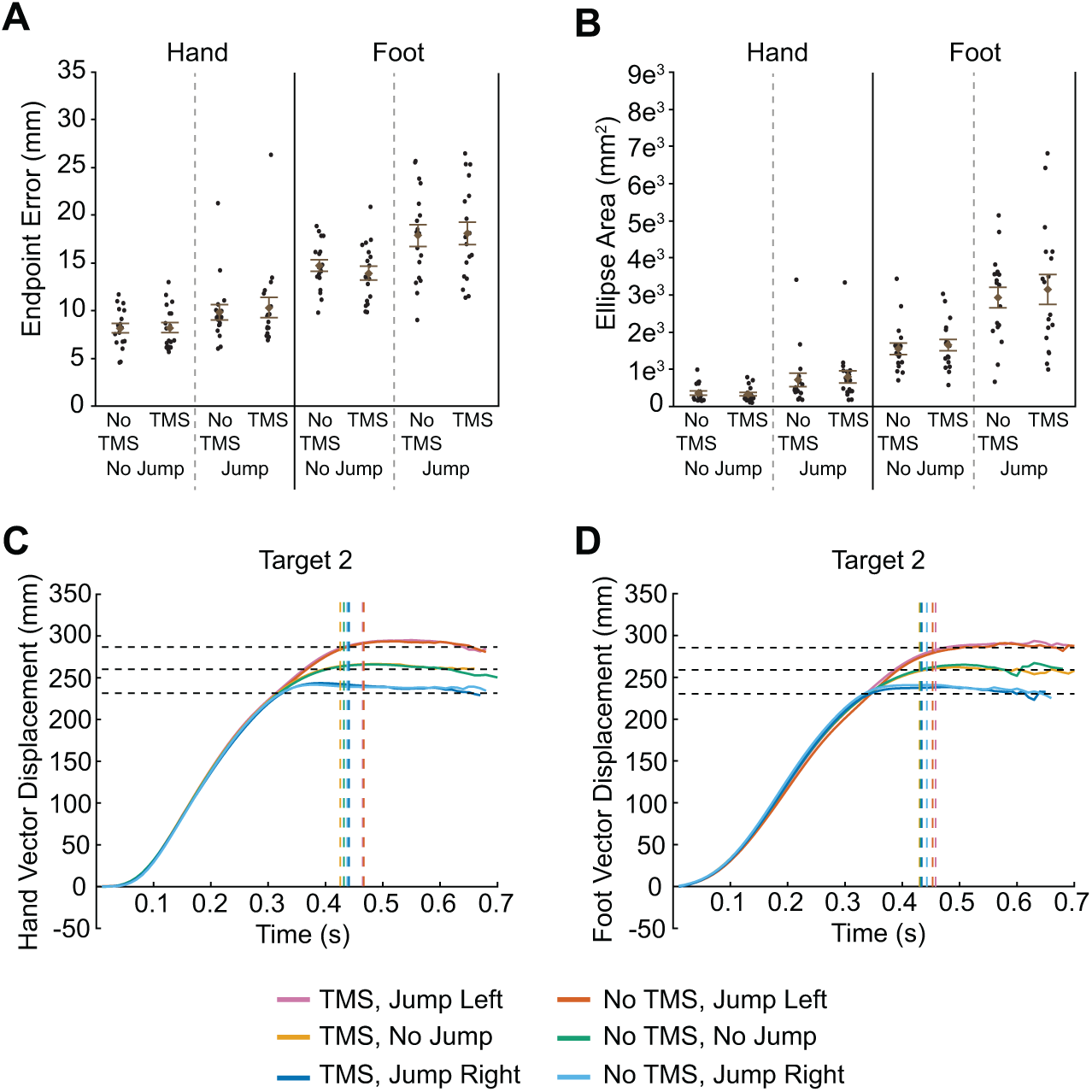
Results of transcranial magnetic stimulation (TMS) to IPS1. Euclidean endpoint error (A) and ellipse area based on endpoint error (B) for hand and foot reaches across all conditions. Filled circles represent mean data from each individual participant. Group mean ± SE is also shown. Hand (C) and foot (D) trajectories towards target two are shown for each condition. Horizontal dashed lines represent the final position of the target (jump left, no jump, and jump right). Vertical dashed lines indicate the average time at which the finger or toe contacted the touchscreen for each condition.

### TMS to IPS1

IPS1 is thought to contribute to both hand and foot motor planning [4,5]. Thus, we predicted that TMS applied to this region would disrupt online corrections following the target jump for both effectors. Disruption may be evident in increased constant error of the reaching effector from the target, or in higher variability of reach endpoints in jump trials when TMS is applied versus when it is not. To test this prediction, we first analyzed finger endpoint error (Fig 2A). Greater error was evident during target-jump trials compared to non-jump trials (F_1,51_ = 15.7, p = 0.0002). However, we found no significant effect of TMS (main effect: F_1,51_ = 0.3, p = 0.567; Jump x TMS: F_1,51_ = 0.2, p = 0.642). Toe endpoint error also increased for jump trials compared to non-jump trials (Fig 2A; F_1,51_ = 42.0, p < 0.0001). Again, however, we found no significant effect of TMS (main effect: F_1,51_ = 0.2, p = 0.628; Jump x TMS: F_1,51_ = 0.8, p = 0.374).

We next determined the predicted ellipse areas based on the endpoint error, a measure of the participant’s error variability. As illustrated in Fig 2B, error variability of hand reaches was greater with jump trials compared to non-jump trials (F_1,51_ = 102.5, p < 0.0001). TMS had no significant effect on error variability (main effect: F_1,51_ = 0.4, p = 0.549; Jump x TMS: F_1,51_ = 3.1, p = 0.085). This figure also shows that error variability of foot reaches increased with jump trials compared to non-jump trials (F_1,51_ = 92.4, p < 0.0001) but there was no significant effect of TMS (main effect: F_1,51_ = 0.5, p = 0.490; Jump x TMS: F_1,51_ = 0.05, p = 0.825).

Although TMS to IPS1 did not affect endpoint error or error variability for hand and foot reaches, it is possible that it affected limb trajectories. Figure 2C,D illustrates the group mean vector displacement of reaches to target 2, separated by jump direction and TMS condition. Trajectories for the other targets are shown in S1 Fig. Both hand and foot trajectories show clear deviations based on target jump direction. However, TMS had no effect as evidenced by the fact that the profiles for TMS and no TMS conditions overlap almost completely. Given this finding, we did not perform any further analyses on these trajectories. We also analyzed vector velocity and acceleration, as well as medial-lateral displacement, but the profiles are similar in that there were no clear effects of TMS.

### TMS to IPS2

Like IPS1, IPS2 is active during both hand and foot motor planning [4,5,7]. As such, we predicted that TMS applied to this region would disrupt online corrections for both hand and foot movements. We found increased endpoint error with target-jump trials compared to non-jump trials for hand movements (Fig 3A; F_1,51_ = 19.8, p < 0.0001) and for foot movements (F_1,51_= 23.6, p < 0.0001). However, we found no significant effect of TMS related to either hand (main effect: F_1,51_ = 0.6, p = 0.446; Jump x TMS: F_1,51_ = 0.7, p = 0.408) or foot (main effect: F_1,51_ = 0.8, p = 0.374; Jump x TMS: F_1,51_ = 0.02, p = 0.881) movements for endpoint error.

**Fig 3.**
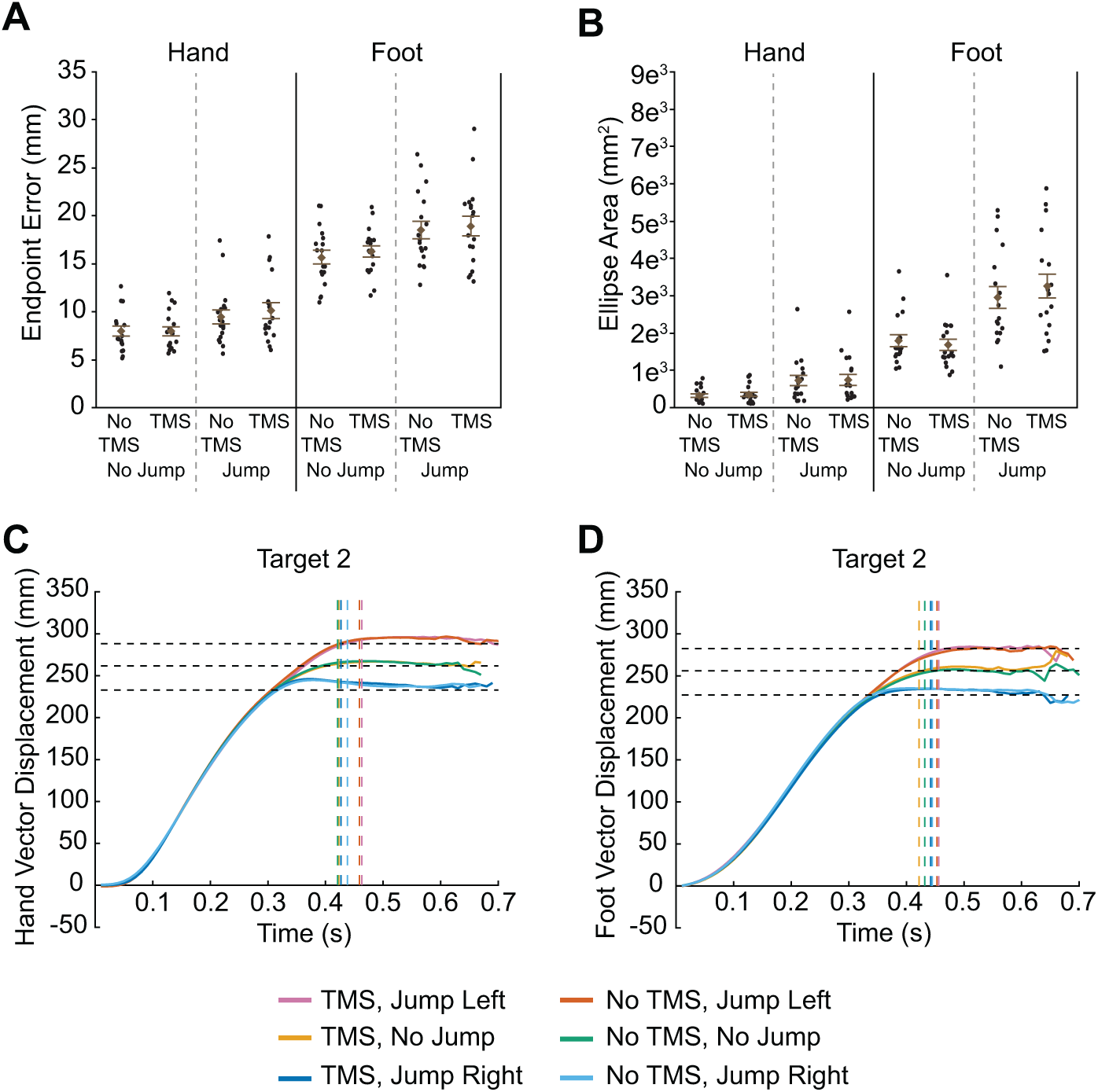
Results of transcranial magnetic stimulation (TMS) to IPS2. Euclidean endpoint error (A) and ellipse area based on endpoint error (B) for hand and foot reaches across all conditions. Filled circles represent mean data from each individual participant. Group mean ± SE is also shown. Hand (C) and foot (D) trajectories towards target two are shown for each condition. Horizontal dashed lines represent the final position of the target (jump left, no jump, and jump right). Vertical dashed lines indicate the average time at which the finger or toe contacted the touchscreen for each condition.

As illustrated in Fig 3B, an analysis on endpoint error variability (i.e., predicted ellipse area) showed greater variability in target-jump trials compared to non-jump trials for hand movements (F_1,51_ = 78.7, p <0.0001) and for foot movements (F_1,51_ = 65.6, p < 0.0001). Once again though, we found no significant effect of TMS related to either hand (main effect: F_1,51_ = 0.2, p = 0.655; Jump x TMS: F_1,51_ = 0.2, p = 0.631) or foot movements (main effect: F_1,51_ = 0.06, p = 0.813; Jump x TMS: F_1,51_ = 1.5, p = 0.232).

Hand and foot trajectories in this TMS condition differed depending on whether the target jumped and which direction it jumped (Fig 3C,D; see also S2 Fig). However, the trajectories with TMS and without TMS for each of the jump directions (no jump, jump left, jump right) were virtually identical, suggesting that TMS had on effect on limb kinematics.

### TMS to handIPS

HandIPS is associated with hand motor planning [33], and activity in this region appears specific to this limb [4,5]. Therefore, we predicted that TMS applied to this brain region would disrupt online corrections to target jumps related to hand movements but not foot movements. Endpoint error (Fig 4A) and predicted ellipse area (Fig 4B) for hand movements is illustrated on the left-hand side of each panel. Similar to the other stimulation conditions discussed above, we found greater hand endpoint error (F_1,51_ = 20.8, p < 0.0001) and ellipse area (F_1,51_ = 89.7, p < 0.0001) for target-jump trials compared to non-jump trials. Contrary to our prediction, however, we found no significant effect of TMS on hand endpoint error (main effect: F_1,51_ = 0.4, p = 0.543; Jump x TMS: F_1,51_ = 0.02, p = 0.888) and ellipse area (main effect: F_1,51_ = 0.05, p = 0.823; Jump x TMS: F_1,51_ = 0.8, p = 0.385).

**Fig 4.**
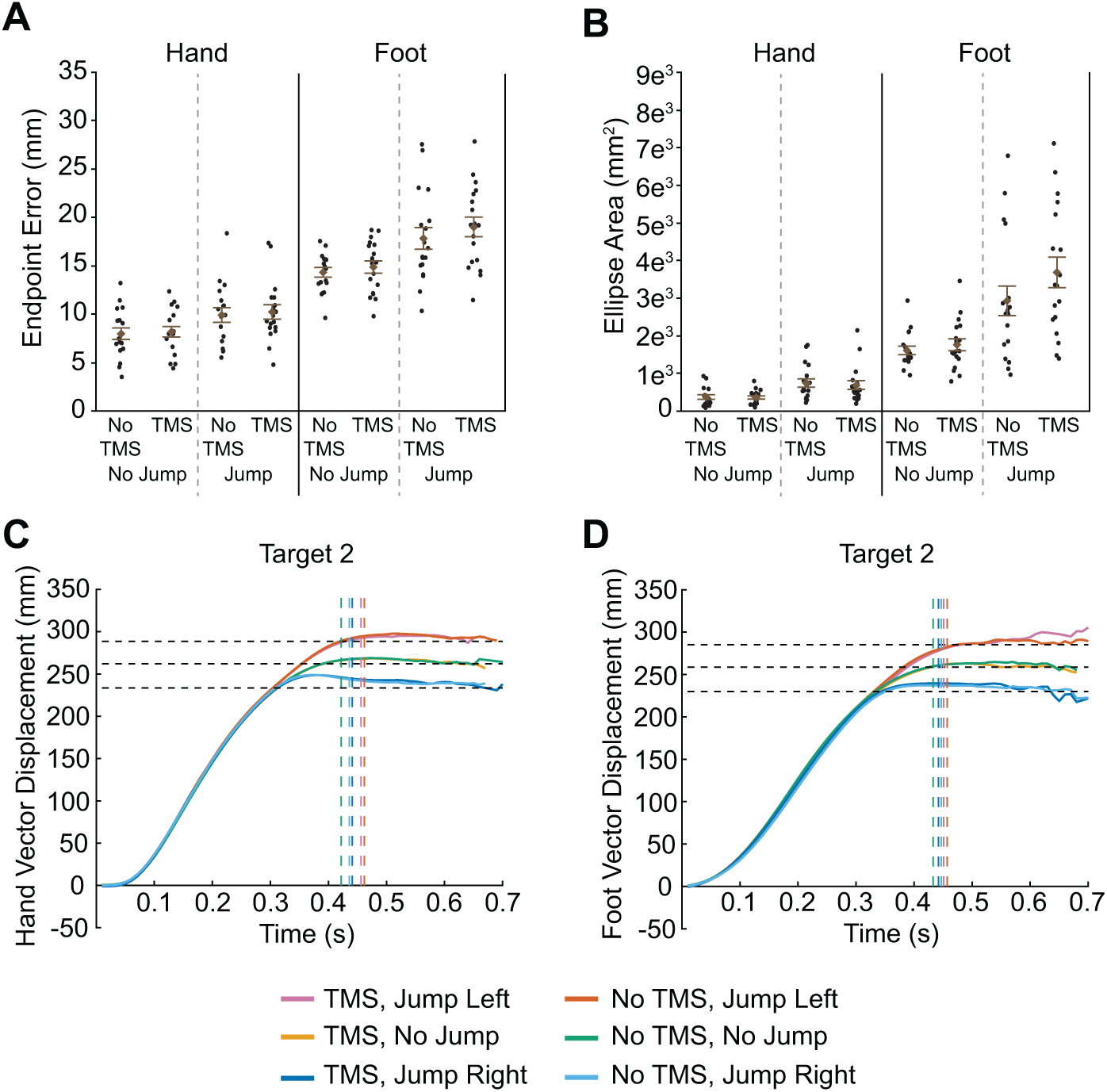
Results of transcranial magnetic stimulation (TMS) to handIPS. Euclidean endpoint error (A) and ellipse area based on endpoint error (B) for hand and foot reaches across all conditions. Filled circles represent mean data from each individual participant. Group mean ± SE is also shown. Hand (C) and foot (D) trajectories towards target two are shown for each condition. Horizontal dashed lines represent the final position of the target (jump left, no jump, and jump right). Vertical dashed lines indicate the average time at which the finger or toe contacted the touchscreen for each condition.

Error data for the foot movements is illustrated on the right-hand side of Fig 4A,B. We found increased foot endpoint error (F_1,51_ = 33.3, p < 0.0001) and ellipse area (F_1,51_ = 57.5, p < 0.0001) for target-jump trials compared to non-jump trials. TMS had no significant effect on foot endpoint error (main effect: F_1,51_ = 1.8, p = 0.189; Jump x TMS: F_1,51_ = 0.3, p = 0.619). Nominally, TMS to handIPS resulted in significantly greater ellipse area for foot endpoint error (main effect: F_1,51_ = 4.4, p = 0.042); note however, that this effect would not survive any correction for multiple testing. The Jump x TMS interaction was not significant (F_1,51_ = 1.7, p = 0.192).

Although TMS to handIPS did not affect endpoint error and ellipse area for hand reaches, it is possible that it disrupted hand trajectories. Figure 4C,D illustrates the group mean vector displacement of both hand and foot reaches to target 2, separated by jump direction and TMS condition. Trajectories for the other targets are shown in S3 Fig. Both hand and foot trajectories show clear deviations based on target jump direction. However, TMS had no effect. This is clearly evident, as the profiles for TMS and no TMS conditions overlap almost completely. Given this finding, we did not perform any further analyses on these trajectories.

### TMS to anterior precuneus

APreC is suspected to contribute to foot-specific motor planning [4,5]. Thus, we predicted that TMS applied to this brain region would disrupt online corrections to target jumps related to foot movements but not hand movements. Figure 5A,B (right-hand side of each panel) illustrates endpoint error and predicted ellipse area for the foot movements to the targets. We found increased endpoint error (F_1,51_ = 25.6, p < 0.0001) and ellipse area (F_1,51_ = 74.6, p < 0.0001) for target-jump trials versus non-jump trials. Although we found no significant effect of TMS on foot endpoint error (main effect: F_1,51_ = 0.2, p = 0.643; Jump x TMS: F_1,51_ = 0.04, p = 0.849), we did find a main effect of TMS for ellipse area (F_1,51_ = 10.1, p = 0.003). Specifically, greater variability was evident with TMS stimulation to aPreC. However, we found no significant interaction for this measure (F_1,51_ = 0.7, p = 0.411) for the foot movement condition.

**Fig 5.**
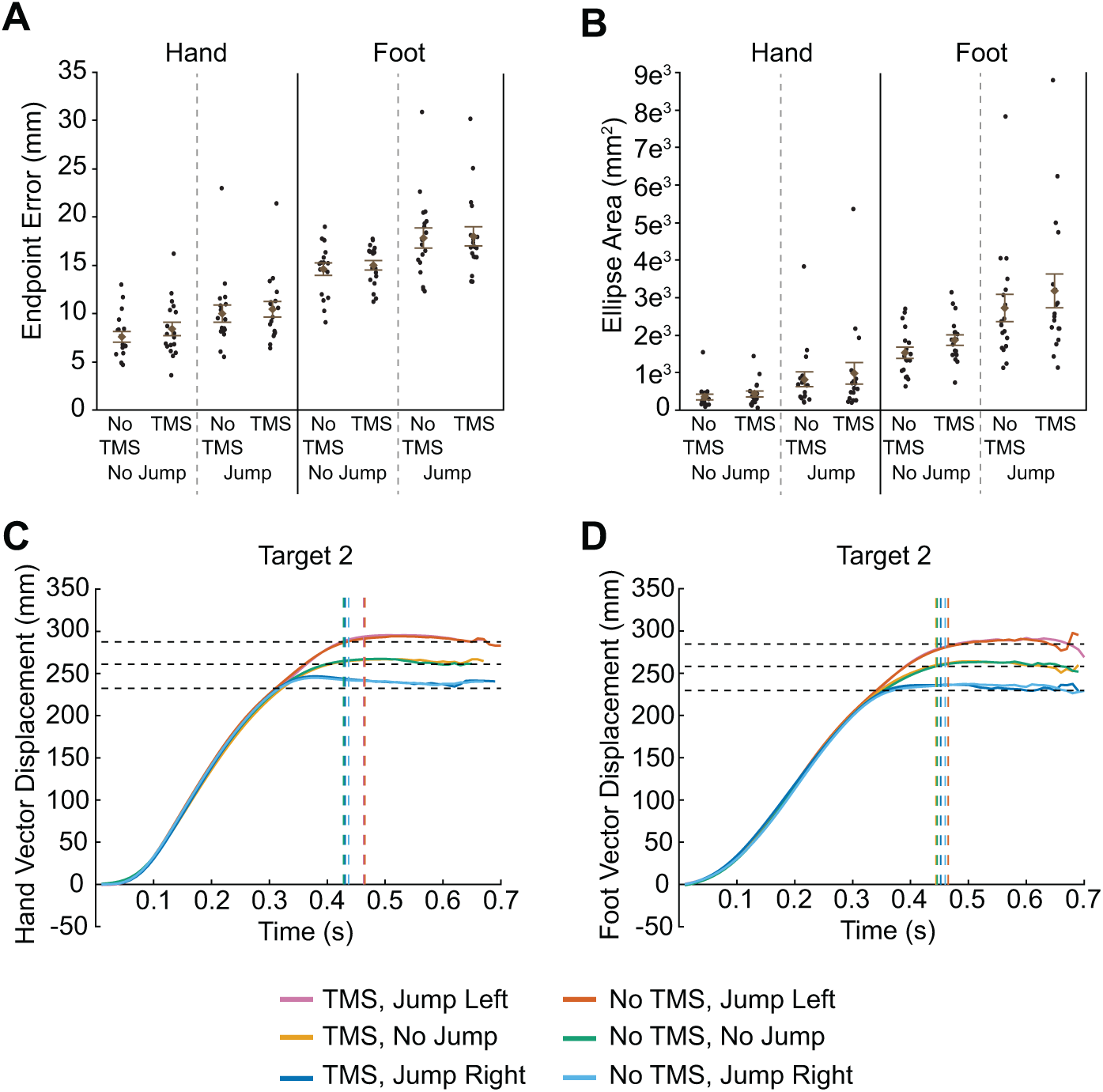
Results of transcranial magnetic stimulation (TMS) to aPreC. Euclidean endpoint error (A) and ellipse area based on endpoint error (B) for hand and foot reaches across all conditions. Filled circles represent mean data from each individual participant. Group mean ± SE is also shown. Hand (C) and foot (D) trajectories towards target two are shown for each condition. Horizontal dashed lines represent the final position of the target (jump left, no jump, and jump right). Vertical dashed lines indicate the average time at which the finger or toe contacted the touchscreen for each condition.

When examining hand movements (Fig 5A,B, left-hand side), we also found increased endpoint error (F_1,51_ = 49.5, p < 0.0001) and ellipse area (F_1,51_ = 68.9, p < 0.0001) for target-jump trials versus non-jump trials. There was a borderline significant effect of TMS on hand endpoint error (F_1,51_ = 4.0, p = 0.050), showing greater error with TMS, but no significant Jump x TMS interaction (F_1,51_ = 0.3, p = 0.574). We also found no significant effect of TMS on ellipse area (main effect: F_1,51_ = 2.3, p = 0.133; Jump x TMS: F_1,51_ = 0.6, p = 0.437).

Hand and foot trajectories in this stimulation condition also differed depending on whether the target jumped to a new location and which direction it jumped (Fig 5C,D; see also S4 Fig). Despite this finding, the trajectories associated with TMS application for each of the jump directions (no jump, jump left, jump right) were virtually identical to those trajectories associated with no TMS application. These results suggest that TMS had no effect on limb kinematics for this stimulation condition.

### Sham TMS

We expected the sham stimulation to have no effect on behaviour. Indeed, sham TMS did not affect endpoint error (Fig 6A). We found no effect of sham TMS for either hand movements (main effect: F_1,51_ = 0.02, p = 0.894; Jump x TMS: F_1,51_ = 0.03, p = 0.860) or foot movements (main effect: F_1,51_ = 0.10, p = 0.718; Jump × TMS: F_1,51_ = 0.1, p = 0.742). Similar to other conditions, endpoint error was greater in target-jump trials versus non-jump trials for hand (F_1,51_ = 42.5, p < 0.0001) and foot movements (F_1,51_ = 28.1, p < 0.0001).

**Fig 6.**
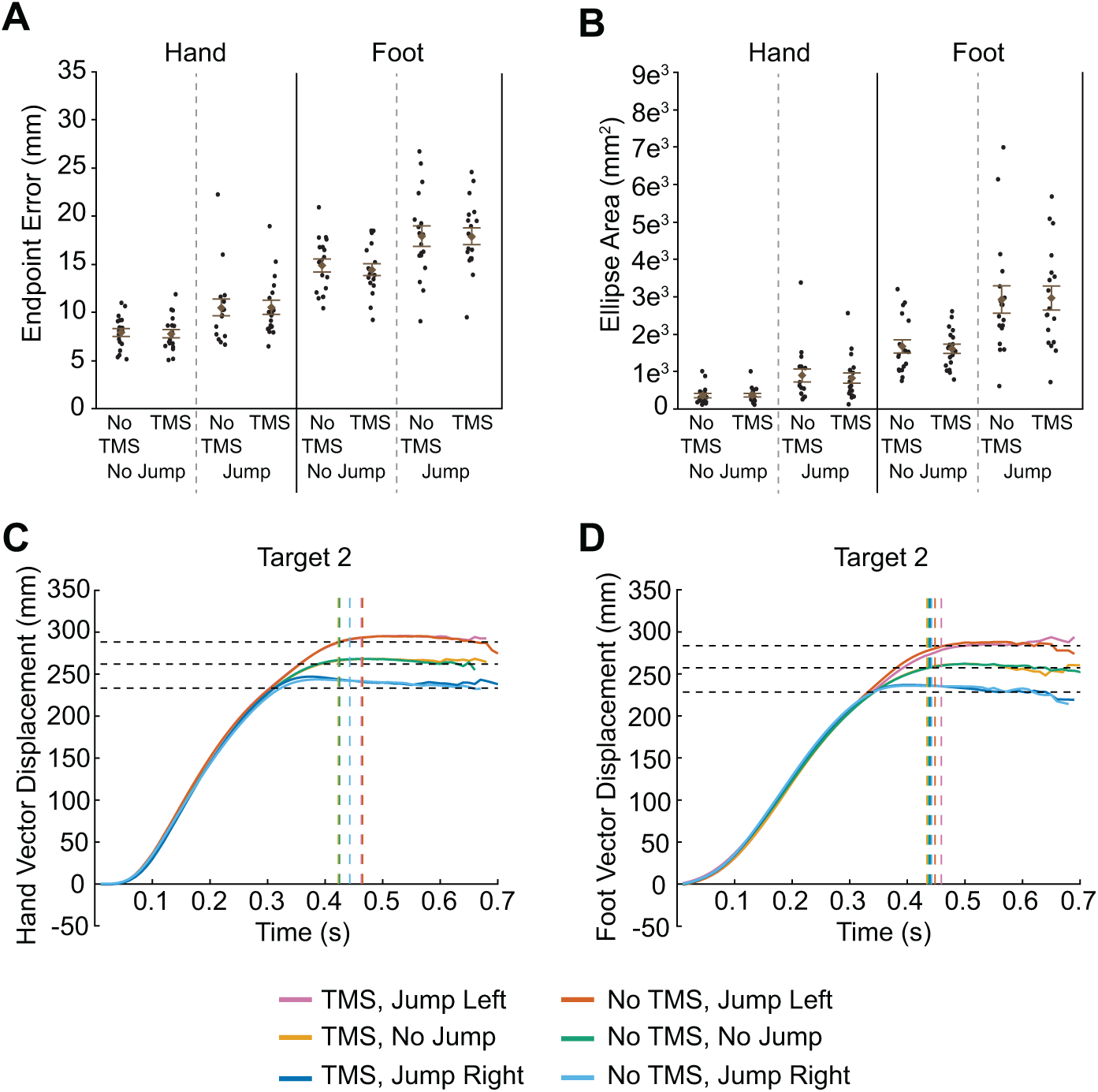
Results of sham stimulation. Euclidean endpoint error (A) and ellipse area based on endpoint error (B) for hand and foot reaches across all conditions. Filled circles represent mean data from each individual participant. Group mean ± SE is also shown. Hand (C) and foot (D) trajectories towards target two are shown for each condition. Horizontal dashed lines represent the final position of the target (jump left, no jump, and jump right). Vertical dashed lines indicate the average time at which the finger or toe contacted the touchscreen for each condition

Next, we examined endpoint error variability, reflected by predicted ellipse area (Fig 6B). Here, we also found no effect of sham TMS for either hand (main effect: F_1,51_ = 0.002, p = 0.969; Jump x TMS: F_1,51_ = 0.5, p = 0.481) or foot (main effect: F_1,51_ = 0.04, p = 0.850; Jump x TMS: F_1,51_ = 0.05, p = 0.831) movements to targets. As expected, error variability was greater in target-jump trials versus non-jump trials for hand (F_1,51_ = 76.1, p < 0.0001) and foot movements (F_1,51_ = 47.7, p < 0.0001).

Finally, we examined limb kinematics. Hand (Fig 6C) and foot trajectories (Fig 6D) differed depending on whether the target jumped and which direction it jumped (see also S5 Fig). However, the trajectories associated with sham TMS for each of the jump directions (no jump, jump left, jump right) were virtually identical to those trajectories associated with no stimulation, suggesting that sham TMS had no effect on limb kinematics.

## Discussion

Human participants executed hand and foot reaches towards visual targets in front of them. In some trials, the target jumped during the movement, requiring participants to adjust their reach online. We attempted to disrupt reach corrections through triple-pulse TMS to one of four parietal regions that are active during the planning and execution of goal-directed hand and foot movements. Here we tested whether the targeted parietal regions are involved in the control of both hand and foot movements, or merely in one or the other. However, we did not observe any variation of movement with TMS, either during regular reaches in which the target remained stable, or during reaches when the target jumped sideways.

Behavioral modulation through TMS can be difficult to achieve, even when targeting a region that is causally involved in the tested behavior. For instance, TMS effects depend on precise targeting, TMS intensity, and sometimes coil orientation. Furthermore, effects can be limited to specific time ranges of TMS relative to experimental events (e.g., [23,39]). Our results may therefore imply that the regions we targeted are not causally involved in hand and foot reaching, but they may alternatively indicate that future research should employ other experimental paradigms that probe different aspects of TMS during reaching, such as the timing and pattern of stimulation. We will discuss these different possible reasons for our negative findings below.

### Consistent TMS targeting

It is possible that we did not correctly position the coil to target the intended brain regions. However, a commercial neuronavigation system, with which others have produced reliable experimental results, guided the TMS, and we based navigation on participants’ individual anatomy assessed with an anatomical MRI scan. Prior to running the study, the experimenters underwent extensive practice in navigating the coil to the correct position and in keeping it on target. We conclude that TMS targeted the regions we intended to stimulate. It is also noteworthy that previous results, acquired both with TMS and fMRI, have resulted in rather variable overall estimates of regions involved in particular aspects of reaching behavior (see, for instance, the review by [2]). In our own previous research, we have found wide-spread activation along both the lateral and medial sides of IPS [4,7]. Thus, slight variation of TMS coil position may be expected to affect some, but probably not all aspects of reaching behavior, and we would have expected TMS effects even if we had employed slightly different TMS coordinates than those defined for the present study..

### Experimental settings: TMS parameters

It is also possible that we did not use optimal stimulation parameters to obtain the desired experimental effects. There is a trade-off in setting stimulation intensity as low as possible so as to minimize the risk of any side effects of TMS for participants, but high enough to obtain sufficient disruption of cortical processing to produce measurable behavioral effects. We determined a resting motor threshold, which is considered less conservative than an active motor threshold, and we used the determined value for experimental TMS. Although different ways of obtaining the motor threshold can render different stimulation intensities, other groups have obtained reliable TMS effects using resting motor threshold and the stimulation intensity we applied here (e.g., [24,36]).

In addition, coil orientation can have an effect when stimulating the motor cortex, and we did not try different orientation configurations. However, we are not aware of any studies that have tested differential effects of coil orientation over the regions we tested here. Therefore, we kept coil orientation equal and in a similar manner to previous studies over all conditions (e.g., [36]).

### Experimental settings: stimulation protocol and timing

The choice of triple-pulse stimulation, too, was motivated by previous reports of successful modulation of reach behavior with multiple short-interval pulses [24,32,36,40]. The underlying idea of this protocol is that several closely timed pulses will affect neural processing more than a single pulse. Furthermore, stimulation covers a larger time interval, so that presumably the specific time point of stimulation is less relevant than the timing of a single pulse. While this methodology, then, renders results that are unspecific regarding the timing of the targeted process, it should be less prone to overlooking a region’s relevance when a stimulation is applied at a non-optimal timepoint. Our three pulses occurred over an 80 ms period of time starting at the target jump. Online corrections to target perturbation are mediated within 110 ms [41]. A previous study applied three TMS pulses from 40 to 73 ms following the perturbation and successfully interrupted online correction [24]. Thus, our stimulation protocol and timing appear adequate to interrupt online correction.

### Choice of regions

We based our choice of regions on three aspects. First, we chose regions that had previously been found active in fMRI experiments during goal-directed hand and/or foot movements. Second, we chose regions that, to our knowledge, had not been disrupted with TMS during foot movement (all regions) or during both hand and foot movement (all regions except IPS1/SPOC). Third, we chose regions that roughly align with the foot regions of M1/S1 and with foot-sensitive neurons in SPL [7,42] on a lateral-to-medial dimension: foot-sensitive neurons are located medially in the M1/S1 homunculus and adjacent SPL. Together, these considerations led us to choose regions with Talairach x-coordinates −26 or closer to zero (left-hemispheric Talairach coordinates are negative, with the medial fissure = 0 and larger |x| indicating more lateral locations). Contrary to the present study, most previous studies have targeted regions located more laterally, usually at Talairach |x| > 35. We briefly discuss each stimulated region, from most posterior to most anterior.

IPS1 is implicated in hand reaching (e.g., [33,43]) and pointing (e.g., [44]; see Figure 2 of [2], for a comprehensive overview). In one study, triple-pulse TMS, the method applied also in the present study, directed to a region about 1 cm medial and posterior to present IPS1, evoked a constant error towards fixation or body midline for hand reaches to lateral targets both with and without vision of the hand [32]; notably, in contrast to the present study, TMS was applied during a planning interval between target presentation and movement execution, rather than online during the movement. Medial reach-related regions were shown to respond to visual target location rather than motor direction when vision was reversed through prisms [45], suggesting that this region codes the target rather than the movement. Accordingly, the vision-independent effect of TMS to IPS1 was interpreted as indicating a disturbance of target processing [32] (with IPS1 referred to as SPOC in their study). Under this notion, both hand- and foot-related reaches in our study should have been sensitive to IPS1 stimulation in the present study, as both relied on identical visual targets.

Our data provide no evidence for an increase of reach error for either limb after stimulating any of our four PPC locations medial to IPS. Previous studies that have interrupted automatic responses to target perturbations have, instead, targeted regions lateral to IPS [13,23,46]. However, we reasoned that an effector-independent effect on reaching should result from TMS during target jumps if, indeed, IPS1 processes the target stimulus rather than the motor response. Given that we applied TMS during movement in our study, the fact that we did not observe an effect of TMS, thus, suggests that IPS1 may be involved in planning, but less, or not at all, in online control of goal-directed limb movements.

To our knowledge, others have not targeted the region we termed IPS2 with TMS during reaching. However, IPS1 and IPS2 are difficult to delineate without dedicated mapping techniques (e.g., [26,27]), and some fMRI activations obtained during reaching [27,31] that have sometimes been subsumed as SPOC/IPS1 [2] are close to the region we refer to as IPS2 here. We included IPS2 in the present study to achieve reasonable coverage of potentially relevant PPC locations along the posterior-anterior axis of PPC. The IPS2 region we chose to stimulate in the present study was active during hand and foot motor planning in previous work, just like IPS1/SPOC [4,7]. Therefore, we expected that TMS to this region during target jumps would affect reach corrections for both limbs. The fact that we did not observe such an effect suggests that, like IPS1, IPS2 may be more planning than execution-related.

Work using fMRI, both during visually and proprioceptively guided pointing and reaching, has implicated the third target region of our study, handIPS, in hand reaching [26,43,47,48]. In fact, this target region is the medial end of a large swath of cortex surrounding the medial IPS, which is involved in mediating both saccades and hand reaches (see [2], for an overview). Notably, this region was not involved in foot movement planning in our previous fMRI studies [4,5,7], and we therefore expected hand-specific TMS effects for this target area. We are not aware of any studies that have aimed as medially as we did here with TMS during reaching. Yet, stimulation in the vicinity, about 1 cm more lateral, affected force judgments while leaving spontaneously produced finger/hand movements intact during probing of artificial force fields [49], in line with our negative results concerning modulation of movement parameters. However, a region 1 cm more lateral and anterior to our handIPS interrupted reaches to visual targets contralateral to stimulation [39], though only if stimulation preceded movement initiation by 100-160 ms. Furthermore, TMS over two comparably lateral but slightly more posterior regions did not disrupt modulation of online correction [24] (their region “Reach” and “SPL”). These latter two regions, too, were within 1-1.5 cm of the handIPS region targeted in the present study. Together with the present result, these findings, thus, suggests that the handIPS region’s function is not related to online control [24].

Finally, the fourth region we stimulated, aPreC, is implicated in hand reaching [30] and pointing [44], as well as in leg and foot movements [4,5,7] (region pCi in [42]; also see [6]). However, in our previous work that used a goal-directed foot pointing task, this region was more active during foot than during hand movement planning [4,5,7].

The aPreC region borders on the M1/S1 foot region; in a similar way, the lateral anterior region over which TMS interrupted online control borders the M1/S1 hand area. The cortex posterior to S1, area 5 or anterior SPL, is organized in a homuncular fashion, similar to S1 [50]. Therefore, we expected that TMS over aPreC would result in disturbance of motor corrections during foot movements. However, because this region was active also during hand movements in some previous studies, we suspected that TMS may also affect hand movements, though presumably less pronounced than foot movements. Such hand and foot-related effects would be consistent with reports of single neuron responses to tactile stimulation of both types of limbs [51,52], even if we targeted an area that was more medial than an SPL region that responded to both hand and foot planning in humans [7]. However, also for this region we did not observe the expected effects of TMS. One possible reason is that we did not stimulate the optimal location within aPreC, so it was not effectively modulated by TMS. However, our previous work identified a large area around the used Talairach coordinate as active during foot movement planning and execution, and so TMS must have affected parts of this region. It is therefore unclear why we did not at least disrupt foot movements when the target jumped during movement. One possibility is that online corrections are not mediated by limb-specific regions that align with the limb-specific sections of primary somatosensory cortex, and thus, foot corrections do not rely on medial SPL. Specifically, the lateral IPS region previously targeted by others to modulate hand online correction [13,23] may be a specialized region for online correction for all limbs, not only the hand. It is noteworthy that, although this more lateral region appeared to be hand-specific in our own previous fMRI studies [4,7] (see also Fig 3 of [5]), in other studies it was active for finger, elbow, and ankle movements (Fig 3 and 5 of [6]) and for hand and foot movement (Fig 3 and 4 of [42]). In fact, others have suggested that this part of PPC may be responsible for online correction independent of the specific limb, based on TMS effects involving corrections with both fingers and wrist [53], as well as single neuron responses related to both hand and shoulder in a human patient with an implant in anterior IPS [54]. Whether such common responses, however, generalize from the hand/arm to the foot/leg also for online control remains an open question.

## Conclusion

In sum, it does not seem plausible to us that the lack of effects in the present study was due merely to the protocol or TMS parameters, although this remains a possibility and will require further exploration. It seems that the regions medial to IPS that we targeted are less involved in online monitoring of movement than assessing the target and motor planning. The present experiment points to two hypotheses that future research can test: first, regions lateral of IPS may be specialized for online control for all limbs and body movements even beyond the hand/arm, much in contrast to the idea that such functionality would be limb-specific and localized near each respective limb’s M1/S1 region. Second, the very medial regions allegedly involved in hand and foot reaching may have other functional roles than online control; in particular, they may be related to movement planning and processing of the movement target. Exploration of these hypotheses will likely require modifications of the methodology applied here, for instance the use of fMRI localizers [24] during perturbations of foot reaches and TMS stimulation at intervals spaced more closely in time than used here [40].

## Acknowledgements

The authors wish to thank Christopher Lau and Liesa Zwetzschke for their help with data collection and participant recruitment.

## References

1. Andersen RA, Cui H. Intention, action planning, and decision making in parietal-frontal circuits. Neuron. 2009;63: 568–583.

2. Vesia M, Crawford JD. Specialization of reach function in human posterior parietal cortex. Exp Brain Res. 2012;221: 1–18.

3. Filimon F. Human cortical control of hand movements: parietofrontal networks for reaching, grasping, and pointing. Neuroscientist. 2010;16: 388–407.

4. Heed T, Beurze SM, Toni I, Röder B, Medendorp WP. Functional rather than effector-specific organization of human posterior parietal cortex. J Neurosci. 2011;31: 3066–3076.

5. Leoné FTM, Heed T, Toni I, Medendorp WP. Understanding effector selectivity in human posterior parietal cortex by combining information patterns and activation measures. J Neurosci. 2014;34: 7102–7112.

6. Cunningham DA, Machado A, Yue GH, Carey JR, Plow EB. Functional somatotopy revealed across multiple cortical regions using a model of complex motor task. Brain Res. 2013;1531: 25–36.

7. Heed T, Leone FTM, Toni I, Medendorp WP. Functional versus effector-specific organization of the human posterior parietal cortex: revisited. J Neurophysiol. 2016;116: 1885–1899.

8. Rijntjes M, Dettmers C, Büchel C, Kiebel S, Frackowiak RSJ, Weiller C. A blueprint for movement: functional and anatomical representations in the human motor system. J Neurosci. 1999;19: 8043–8048.

9. Miall RC, Wolpert DM. Forward models for physiological motor control. Neural Net. 1996;9: 1265–1279.

10. Mulliken GH, Musallam S, Andersen RA. Forward estimation of movement state in posterior parietal cortex. Proc Natl Acad Sci USA. 2008;105: 8170–8177.

11. Wolpert DM, Goodbody SJ, Husain M. Maintaining internal representations: the role of the human superior parietal lobe. Nat Neurosci. 1998;1: 529–533.

12. Buneo CA, Andersen RA. The posterior parietal cortex: sensorimotor interface for the planning and online control of visually guided movements. Neuropsychologia. 2006; 44: 2594–2606.

13. Desmurget M, Epstein CM, Turner RS, Prablanc C, Alexander GE, Grafton ST. Role of the posterior parietal cortex in updating reaching movements to a visual target. Nat Neurosci. 1999;2: 563–567.

14. Drew T, Marigold DS. Taking the next step: cortical contributions to the control of locomotion. Curr Opin Neurobiol. 2015;33: 25–33.

15. Marigold DS, Drew T. Posterior parietal cortex estimates the relationship between object and body location during locomotion. eLife. 2017;6:e28143.

16. Marigold DS, Andujar J-E, Lajoie K, Drew T. Motor planning of locomotor adaptations on the basis of vision: the role of the posterior parietal cortex. Prog Brain Res. 2011;188: 83–100.

17. Danckert J, Ferber S, Goodale MA. Direct effects of prismatic lenses on visuomotor control: an event-related functional MRI study. Eur J Neurosci. 2008;28: 1696–1704.

18. Luauté J, Schwartz S, Rossetti Y, Spiridon M, Rode G, Boisson D, et al. Dynamic changes in brain activity during prism adaptation. J Neurosci. 2009;29: 169–178.

19. Archambault PS, Caminiti R, Battaglia-Mayer A. Cortical mechanisms for online control of hand movement trajectory: the role of the posterior parietal cortex. Cereb Cortex. 2009;19: 2848–2864.

20. Archambault PS, Ferrari-Toniolo S, Battaglia-Mayer A. Online control of hand trajectory and evolution of motor intention in the parietofrontal system. J Neurosci. 2011;31: 742–752.

21. Marigold DS, Drew T. Contribution of cells in the posterior parietal cortex to the planning of visually guided locomotion in the cat: effects of temporary visual interruption. J Neurophysiol. 2011;105: 2457–2470.

22. Gréa H, Pisella L, Rossetti Y, Desmurget M, Tilikete C, Grafton S, et al. A lesion of the posterior parietal cortex disrupts on-line adjustments during aiming movements. Neuropsychologia. 2002;40: 2471–2480.

23. Tunik E, Frey SH, Grafton ST. Virtual lesions of the anterior intraparietal area disrupt goal-dependent on-line adjustments of grasp. Nat Neurosci. 2005;8: 505–511.

24. Reichenbach A, Bresciani J-P, Peer A, Bülthoff HH, Thielscher A. Contributions of the PPC to online control of visually guided reaching movements assessed with fMRI-guided TMS. Cereb Cortex. 2011;21: 1602–1612.

25. Sandrini M, Umiltà C, Rusconi E. The use of transcranial magnetic stimulation in cognitive neuroscience: A new synthesis of methodological issues. Neurosci Biobehav Rev. 2011;35: 516–536.

26. Hagler DJ Jr, Riecke L, Sereno MI. Parietal and superior frontal visuospatial maps activated by pointing and saccades. NeuroImage. 2007;35: 1562–1577.

27. Levy I, Schluppeck D, Heeger DJ, Glimcher PW. Specificity of human cortical areas for reaches and saccades. J Neurosci. 2007;27: 4687–4696.

28. Schluppeck D, Glimcher P, Heeger DJ. Topographic organization for delayed saccades in human posterior parietal cortex. J Neurophysiol. 2005;94: 1372–1384.

29. Silver MA, Ress D, Heeger DJ. Topographic maps of visual spatial attention in human parietal cortex. J Neurophysiol. 2005;94: 1358–1371.

30. Filimon F, Nelson JD, Huang R-S, Sereno MI. Multiple parietal reach regions in humans: cortical representations for visual and proprioceptive feedback during on-line reaching. J Neurosci. 2009;29: 2961–2971.

31. Prado J, Clavagnier S, Otzenberger H, Scheiber C, Kennedy H, Perenin MT. Two cortical systems for reaching in central and peripheral vision. Neuron. 2005;48: 849–858.

32. Vesia M, Prime SL, Yan X, Sergio LE, Crawford JD. Specificity of human parietal saccade and reach regions during transcranial magnetic stimulation. J Neurosci. 2010;30: 13053–13065.

33. Blangero A, Menz MM, McNamara A, Binkofski F. Parietal modules for reaching. Neuropsychologia. 2009;47: 1500–1507.

34. Glover S, Miall RC, Rushworth MFS. Parietal rTMS disrupts the initiation but not the execution of on-line adjustments to a perturbation of object size. J Cogn Neurosci. 2005;17: 124–136.

35. Le A, Vesia M, Yan X, Crawford JD, Niemeier M. Parietal area BA7 integrates motor programs for reaching, grasping, and bimanual coordination. J Neurophysiol. 2017;117: 624–636.

36. Striemer CL, Chouinard PA, Goodale MA. Programs for action in superior parietal cortex: a triple-pulse TMS investigation. Neuropsychologia. 2011;49: 2391–2399.

37. Rossi S, Hallett M, Rossini PM, Pascual-Leone A, the Safety of TMS Consensus Group. Safety, ethical considerations, and application guidelines for the use of transcranial magnetic stimulation in clinical practice and research. Clin Neurophysiol. 2009;120: 2008–2039.

38. Schubert P, Kirchner M. Ellipse area calculations and their applicability in posturography. Gait Posture. 2014;39: 518–522.

39. Davare M, Zénon A, Desmurget M, Olivier E. Dissociable contribution of the parietal and frontal cortex to coding movement direction and amplitude. Front Hum Neurosci. 2015;9.

40. Reichenbach A, Bresciani J-P, Bülthoff HH, Thielscher A. Reaching with the sixth sense: vestibular contributions to voluntary motor control in the human right parietal cortex. NeuroImage. 2016;124: 869–875.

41. Scott SH. A Functional taxonomy of bottom-up sensory feedback processing for motor actions. Trends Neurosci. 2016;39: 512–526.

42. Serra C, Galletti C, Marco SD, Fattori P, Galati G, Sulpizio V, et al. Egomotion-related visual areas respond to active leg movements. Hum Brain Mapp. doi: 10.1002/hbm.24589

43. Beurze SM, de Lange FP, Toni I, Medendorp WP. Spatial and effector processing in the human parietofrontal network for reaches and saccades. J Neurophysiol. 2009;101: 3053–3062.

44. Tosoni A, Galati G, Romani GL, Corbetta M. Sensory-motor mechanisms in human parietal cortex underlie arbitrary visual decisions. Nat Neurosci. 2008;11: 1446–1453.

45. Fernandez-Ruiz J, Goltz HC, DeSouza JFX, Vilis T, Crawford JD. Human parietal “reach region” primarily encodes intrinsic visual direction, not extrinsic movement direction, in a visual-motor dissociation task. Cereb Cortex. 2007;17: 2283–2292.

46. Della-Maggiore V, Malfait N, Ostry DJ, Paus T. Stimulation of the posterior parietal cortex interferes with arm trajectory adjustments during the learning of new dynamics. J Neurosci. 2004;24: 9971–9976.

47. Medendorp WP, Goltz HC, Crawford JD, Vilis T. Integration of target and effector information in human posterior parietal cortex for the planning of action. J Neurophysiol. 2005;93: 954–962.

48. Pellijeff A, Bonilha L, Morgan PS, McKenzie K, Jackson SR. Parietal updating of limb posture: an event-related fMRI study. Neuropsychologia. 2006;44: 2685–2690.

49. Leib R, Mawase F, Karniel A, Donchin O, Rothwell J, Nisky I, et al. Stimulation of PPC affects the mapping between motion and force signals for stiffness perception but not motion control. J Neurosci. 2016;36: 10545–10559.

50. Taoka M, Toda T, Iwamura Y. Representation of the midline trunk, bilateral arms, and shoulders in the monkey postcentral somatosensory cortex. Exp Brain Res. 1998;123: 315–322.

51. Breveglieri R, Galletti C, Gamberini M, Passarelli L, Fattori P. Somatosensory cells in area PEc of macaque posterior parietal cortex. J Neurosci. 2006;26: 3679–3684.

52. Breveglieri R, Galletti C, Monaco S, Fattori P. Visual, somatosensory, and bimodal activities in the macaque parietal area PEc. Cereb Cortex. 2008;18, 806–816.

53. Tunik E, Rice NJ, Hamilton A, Grafton ST. Beyond grasping: representation of action in human anterior intraparietal sulcus. NeuroImage. 2007;36: T77–T86.

54. Zhang CY, Aflalo T, Revechkis B, Rosario ER, Ouellette D, Pouratian N, et al. Partially mixed selectivity in human posterior parietal association cortex. Neuron. 2017;95, 697–708.

