## Supplementary Figures for "No effect of triple-pulse TMS medial to intraparietal sulcus on online correction for target perturbations during goal-directed hand and foot reaches"

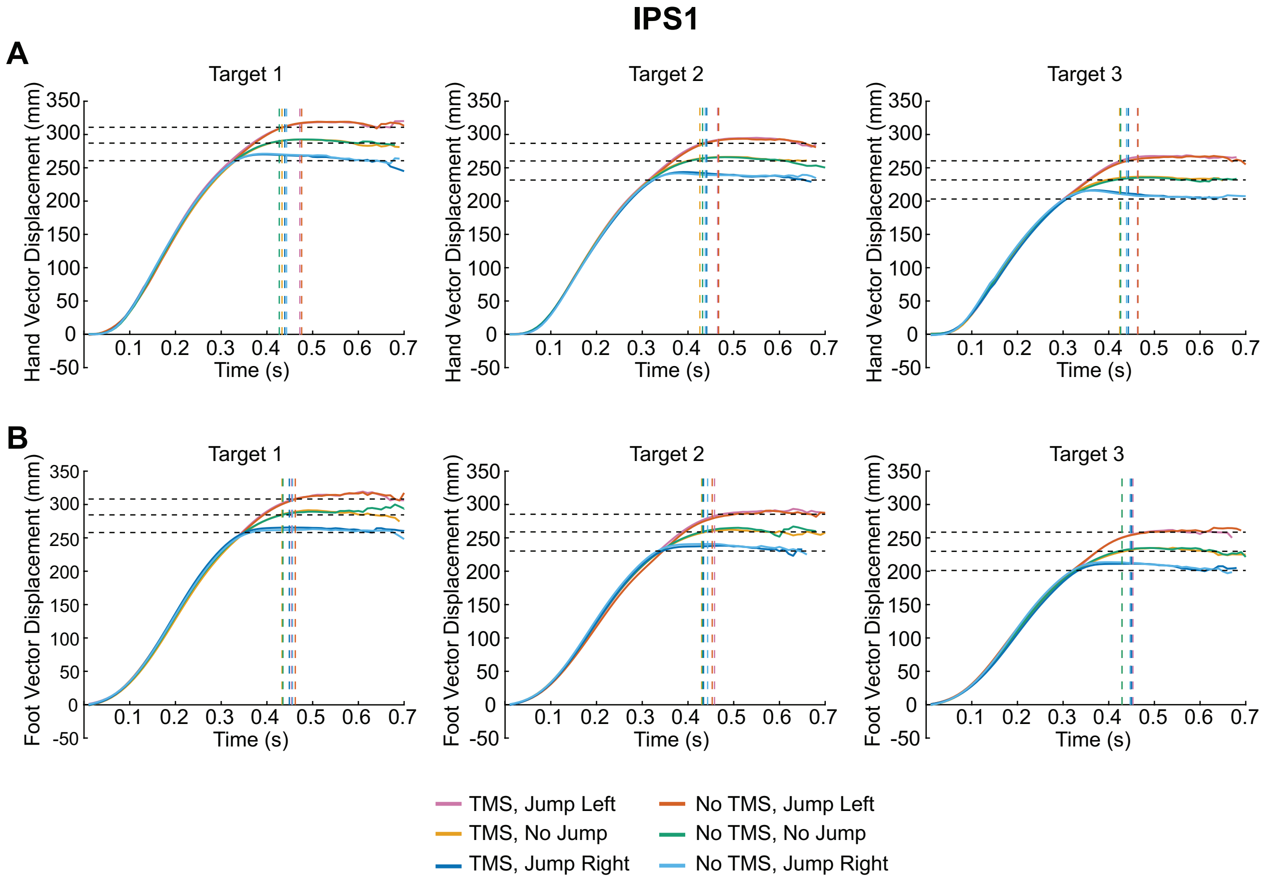


### S1 Fig

**Effect of transcranial magnetic stimulation (TMS) to IPS1 on reach trajectories.** Group mean hand (A) and foot (B) trajectories towards targets are shown for each condition. Horizontal dashed lines represent the final position of the target (jump left, no jump, and jump right). Vertical dashed lines indicate the average time at which the finger or toe contacted the touchscreen for each condition.

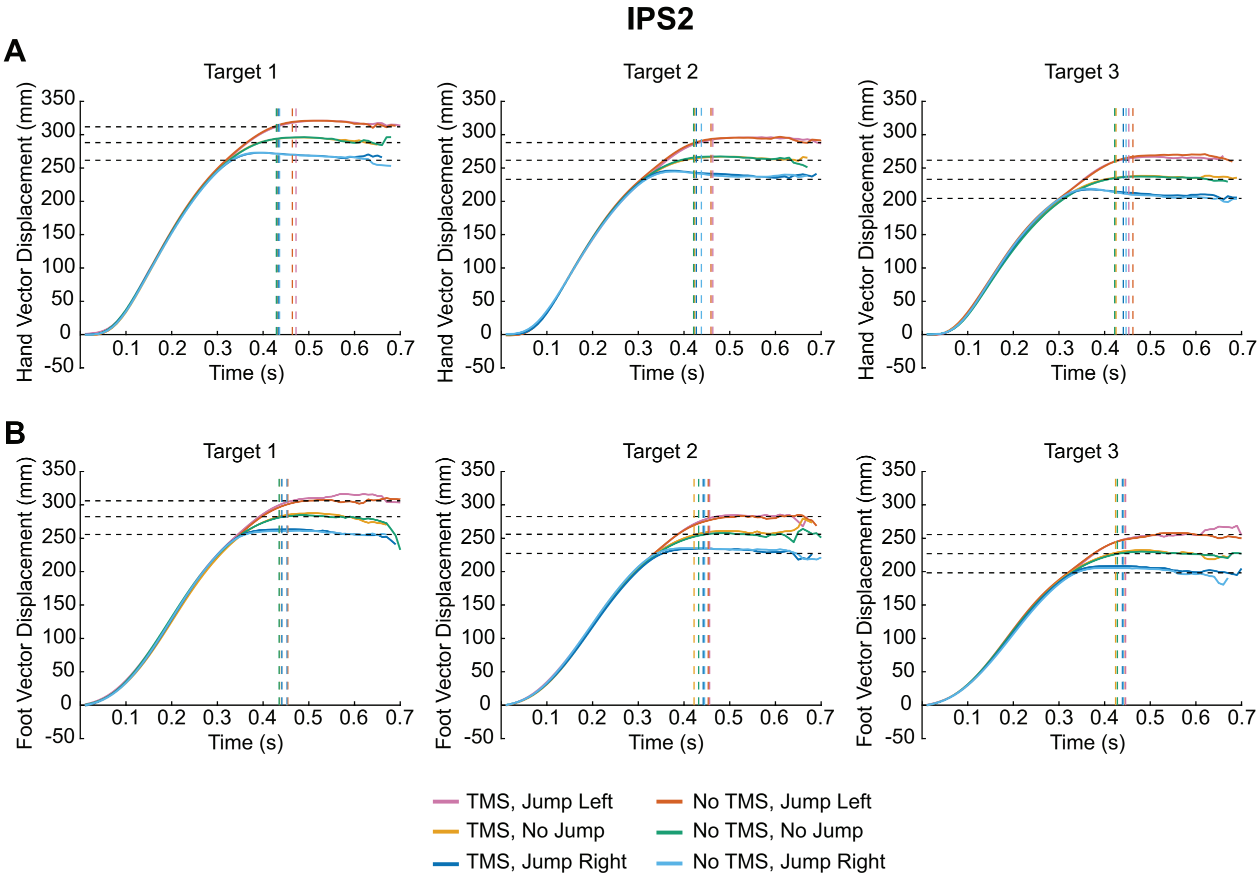


### S2 Fig

**Effect of transcranial magnetic stimulation (TMS) to IPS2 on reach trajectories.** Group mean hand (A) and foot (B) trajectories towards targets are shown for each condition. Horizontal dashed lines represent the final position of the target (jump left, no jump, and jump right). Vertical dashed lines indicate the average time at which the finger or toe contacted the touchscreen for each condition.

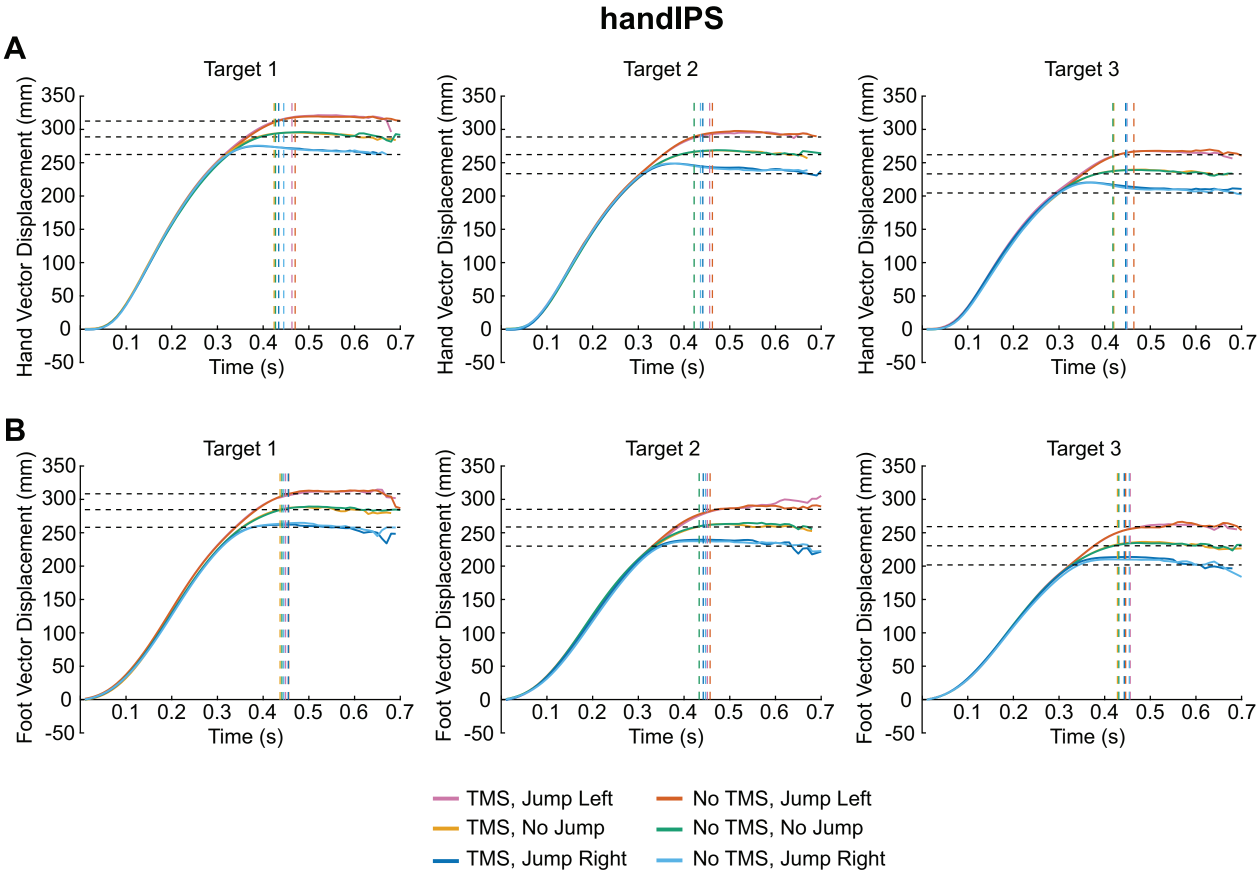


### S3 Fig

**Effect of transcranial magnetic stimulation (TMS) to handIPS on reach trajectories.** Group mean hand (A) and foot (B) trajectories towards targets are shown for each condition. Horizontal dashed lines represent the final position of the target (jump left, no jump, and jump right). Vertical dashed lines indicate the average time at which the finger or toe contacted the touchscreen for each condition.

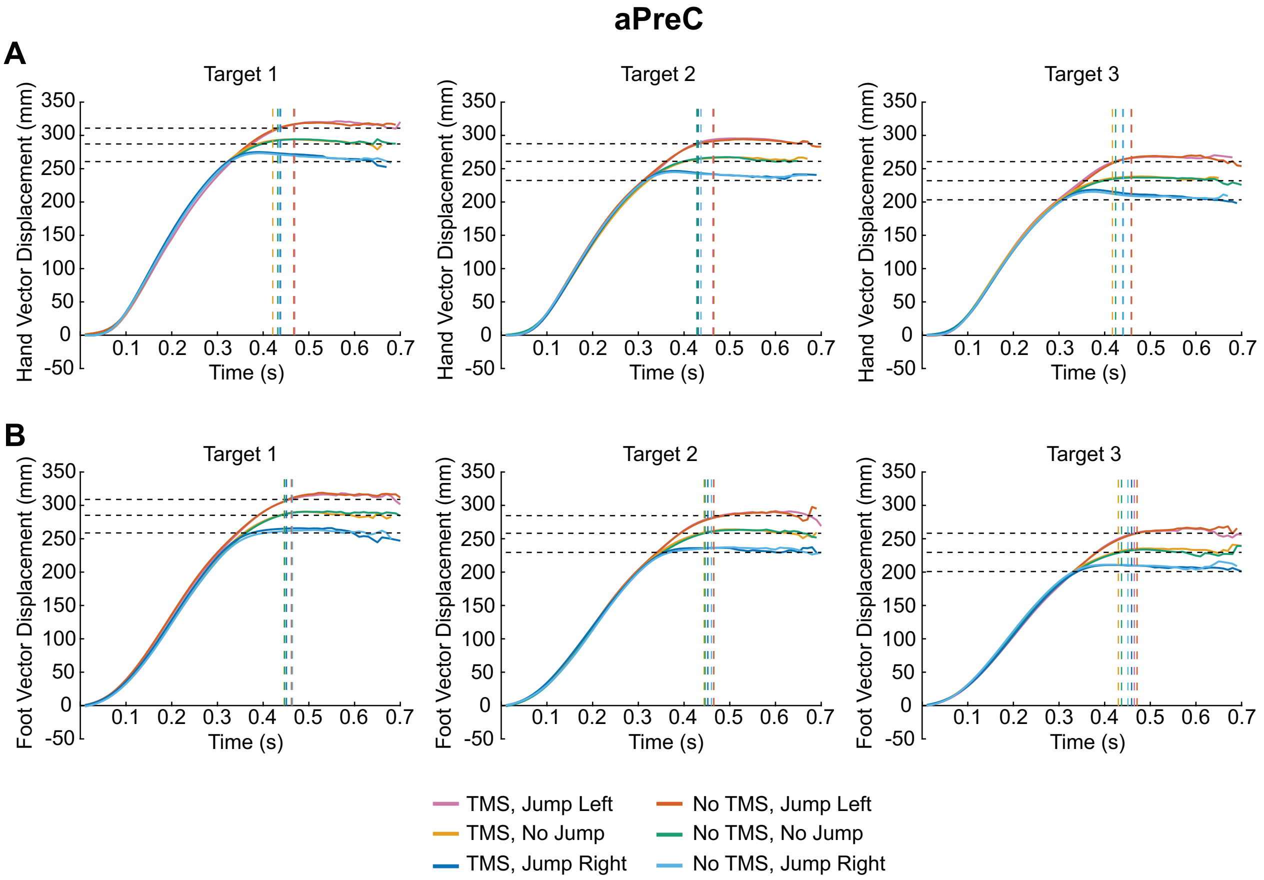


### S4 Fig

**Effect of transcranial magnetic stimulation (TMS) to aPreC on reach trajectories.** Group mean hand (A) and foot (B) trajectories towards targets are shown for each condition. Horizontal dashed lines represent the final position of the target (jump left, no jump, and jump right). Vertical dashed lines indicate the average time at which the finger or toe contacted the touchscreen for each condition.

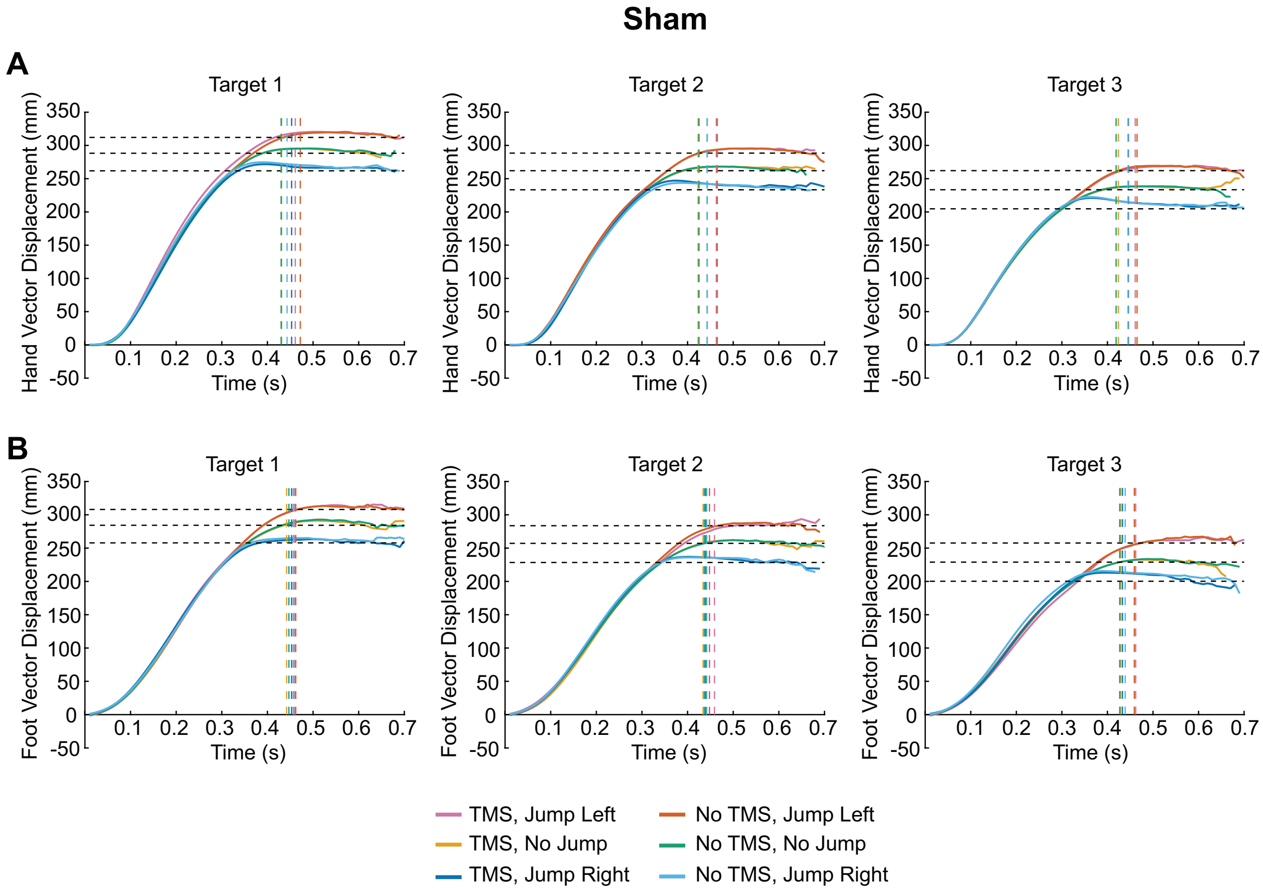


### S5 Fig

**Effect of sham transcranial magnetic stimulation (TMS) on reach trajectories.** Group mean hand (A) and foot (B) trajectories towards targets are shown for each condition. Horizontal dashed lines represent the final position of the target (jump left, no jump, and jump right). Vertical dashed lines indicate the average time at which the finger or toe contacted the touchscreen for each condition.
